# Effect sizes and standardization in neighborhood models of forest stands

**DOI:** 10.1101/010520

**Authors:** Peter Stoll, David J. Murrell, David M. Newbery

## Abstract

Effects of conspecific neighbors on growth and survival of trees have been found to be related to species abundance. Both positive and negative relationships may explain observed abundance patterns. Surprisingly, it is rarely tested whether such relationships could be biased or even spurious due to influences of spatial aggregation, distance decay of neighbor effects and standardization of effect sizes. To investigate potential biases, we simulated communities of 20 identical species with log-series abundances but without species-specific interactions. We expected no relationship of conspecific neighbor effects with species abundance. Growth of individual trees was simulated in random and aggregated spatial patterns using no, linear, or squared distance decay. Regression coefficients of statistical neighborhood models were unbiased and unrelated to species abundance. However, variation in the number of conspecific neighbors was positively or negatively related to species abundance depending on spatial pattern and type of distance decay. Consequently, effect sizes and standardized regression coefficients were also positively or negatively related to species abundance depending on spatial pattern and distance decay. We argue that tests using randomized tree positions and identities provide the best bench marks by which to critically evaluate relationships of effect sizes or standardized regression coefficients with tree species abundance.

## 1 Introduction

Whether or not conspecific negative density dependence (CNDD) at small neighborhood scales shapes species abundances in tropical tree communities at larger scales is far from resolved and we probably should not even expect the answer to be simple. In principle, there are several possibilities. First, the strength of CNDD is unrelated to abundance. Second, the strength of CNDD is negatively related to abundance (strong CNDD for abundant but weak for rare species). This would prevent abundant species to become even more abundant and competitively exclude other species. Moreover, it would confer a rare-species advantage and possibly lead to a community compensatory trend (CCT, Connell et al. 1984). Third, the strength of CNDD is positively related to abundance (strong CNDD for rare but weak for abundant species). This would explain the rarity and low abundance of the species with strong CNDD and the high abundances of species with weak CNDD (Comita et al. 2010). There remain, though, two further possibilities, *viz*. that a mix of positive and negative processes is operating, or the observed relationships are simply spurious (i.e. the result of a statistical artefact).

Recently published experimental results showed that negative density dependence caused by fungal pathogens and insect herbivores was greatest for the species that were most abundant as seeds (Bagchi et al. 2014). In contrast, positive relationships between a species’ average abundance and its negative density dependence (Comita et al. 2010) could be explained if the causality is reversed and lower negative density dependence leads to increased abundance. Moreover, small seeds are likely to be more vulnerable to natural enemies, and small seeds are produced in greater numbers. Thus, differences in seed size may reconcile Bagchi and colleagues’ results with previous work (Muller-Landau 2014).

We investigated relationships between the strength of CNDD and abundance using a simple, spatially explicit and individual-based model simulating identical species without any species-specific interactions. Thus, we would not expect any relationships between the strength of CNDD and abundance in communities simulated under these assumptions. Nevertheless, relationships do emerge because of interfering effects of spatial patterns or distance decay (i.e. the functional form relating neighbor effects to distance from focal trees, Fig. 1) and, perhaps most importantly, due to the common practice of scaling input variables. For example, if rare species have higher variance in the number of conspecifics in their local neighborhoods compared to common species, scaling is expected to increase effect sizes (or standardized partial correlation coefficients) of rare relative to common species, possibly leading to spurious positive relations between the strength of CNDD and abundances. Scaling or standardization is usually recommended (e.g., Schielzeth 2010) and applied especially in hierarchical Bayesian modeling to speed up or even ensure numerical convergence (e.g., Gelman and Hill 2007).

**Figure. 1.**
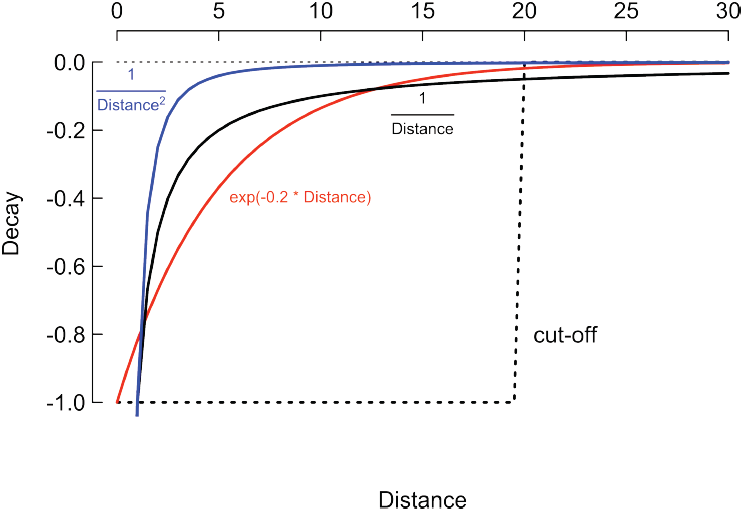
Distance decay of neighborhood effects. In the cut-off model (dashed), the sizes of bigger neighbors with a distance < cut-off are summed. In the linear distance decay (black), the sizes of bigger neighbors are weighed by 1/distance. This is similar to an exponential distance decay (red), which, however, gives somewhat more weight at intermediate distances. A decay of 1/distane^2^ (blue) yields a very rapidly decreasing function. Beyond 20, all three functions give essentially zero weights.

Our initial motivation to investigate relationships between the strength of CNDD and abundance using simulations was two-fold. First, we were puzzled by a consistent negative relationship between the strength of CNDD (i.e. effect sizes derived from statistical neighborhood models) and abundance (total basal area of species) in randomization tests of our own results (Newbery and Stoll 2013). Second, a positive relationship between the strength of CNDD and abundance was found by Comita et al. (2010). Such contrasting results are very interesting if they relate to different underlying biological mechanisms operating on different species in different localities, but we should first try to rule out any differences that might be caused by statistical methods.

## 2 Materials and Methods

We simulated a completely neutral forest without any species-specific effects. Initial size distributions of individuals (basal area, *ba*) were log-normal with mean 2 and standard deviation 1. Individuals of 20 identical species with log-series abundances (i.e. 2827, 1408, 935, 699, 557, 462, 395, 344, 305, 273, 248, 226, 208, 192, 179, 167, 157, 147, 139, 132) were placed on plots (200 x 400 m) either randomly or with aggregated spatial patterns. The aggregated pattern was realized by dispersing individuals around ‘parent trees’ (assigned random locations according to a homogeneous Poisson process), using a Gaussian dispersal kernel with mean 0 and standard deviation 3 m. Thus the species distributions were modeled as a Thomas cluster process, which in turn is a special case of a Neyman-Scott cluster process (Neyman and Scott 1952), and this method means species are spatially independent of one another. For each individual, one single growth increment (absolute growth rate, *agr*) was simulated for trees within a border of 20 m using the following multiple regression equation:

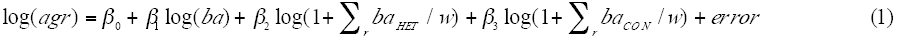

with w = 1 (no distance decay), w = distance (linear distance dacay) or w = distance^2^ (squared distance decay, Fig. 1). The neighborhood terms (ba_HET_ and ba_CON_) summed the basal areas of bigger heterospecific (HET) or bigger conspecific (CON) neighbors within a neighborhood radius (r) of 20 m. The random error term was N (0, 0.3). Regression coefficients were β_0_ = -0.1, β_1_ = 0.3 and β_2_ = β_3_ = -0.2. To verify the simulations, test runs with random errors set to N (0, 0) were performed. The simulations were realized using C++ (computer code is given in Appendix A of the supplementary material).

Neighborhood models (as in Stoll and Newbery 2005) were then fitted to the simulated data over all possible combinations of radii for HET and CON neighbors using R (R Development Core Team 2012) and parameter estimates taken from those models yielding the highest adjusted R^2^-values. Five runs with different seeds were performed and estimates of regression coefficients from best fitting neighborhood models, effect sizes (Cohen 1988; Nakagawa and Cuthill 2007) or standardized regression coefficients (e.g., Warner 2012) averaged across the five runs. Effect sizes (i.e. squared partial correlation coefficients, t^2^ /[t^2^ + residual degrees of freedom], t = t-value) and standardized regression coefficients (b = β’s obtained from regressions with all input variables standardized by subtracting their mean and dividing by their standard deviation) were then correlated with species abundances (i.e. plot level basal area, BA, log transformed). Standardized regression coefficients can be calculated from unstandardized β’s as b = β * SD_X_ / SD_Y_. A positive correlation implies that less abundant, rare species have stronger CON effects – β is more negative – (as in Comita et al. 2010), whereas a negative relationship implies more abundant species have stronger CON effects (as in Newbery and Stoll 2013).

## 3 Results

There were no significant regressions of negative conspecific density dependence (regression coefficient β_3_ in Eq. 1) and species abundance (plot level basal area) regardless of distance decay or spatial pattern (Fig. 2). Variation in parameter estimates was largest for squared distance decay and random spatial pattern. Best fitting radii for bigger conspecific neighbors were unbiased in neighborhood models without distance decay and random spatial pattern (Table 1). However, in the aggregated pattern and with linear distance decay they were slightly underestimated. With estimates (mean ± SD) of 15.9 ± 2.6 in the random spatial pattern and 14.5 ± 3.2, the underestimation was more pronounced with squared distance decay.

**TABLE 1.** Average best fitting radii ± standard deviation (SD) for bigger conspecifics in neighborhood models (Eq. 1) across 20 species with identical initial size distributions and log-species abundances and random or aggregated spatial patterns.

| Distance decay | Spatial pattern |  |  |  |  |  |
| --- | --- | --- | --- | --- | --- | --- |
|  | random |  |  | aggregated |  |  |
| no | 20.0 | ± | 0.0 | 19.8 | ± | 0.4 |
| linear | 19.6 | ± | 0.5 | 19.1 | ± | 1.0 |
| squared | 15.9 | ± | 2.6 | 14.5 | ± | 3.2 |

**Figure. 2.**
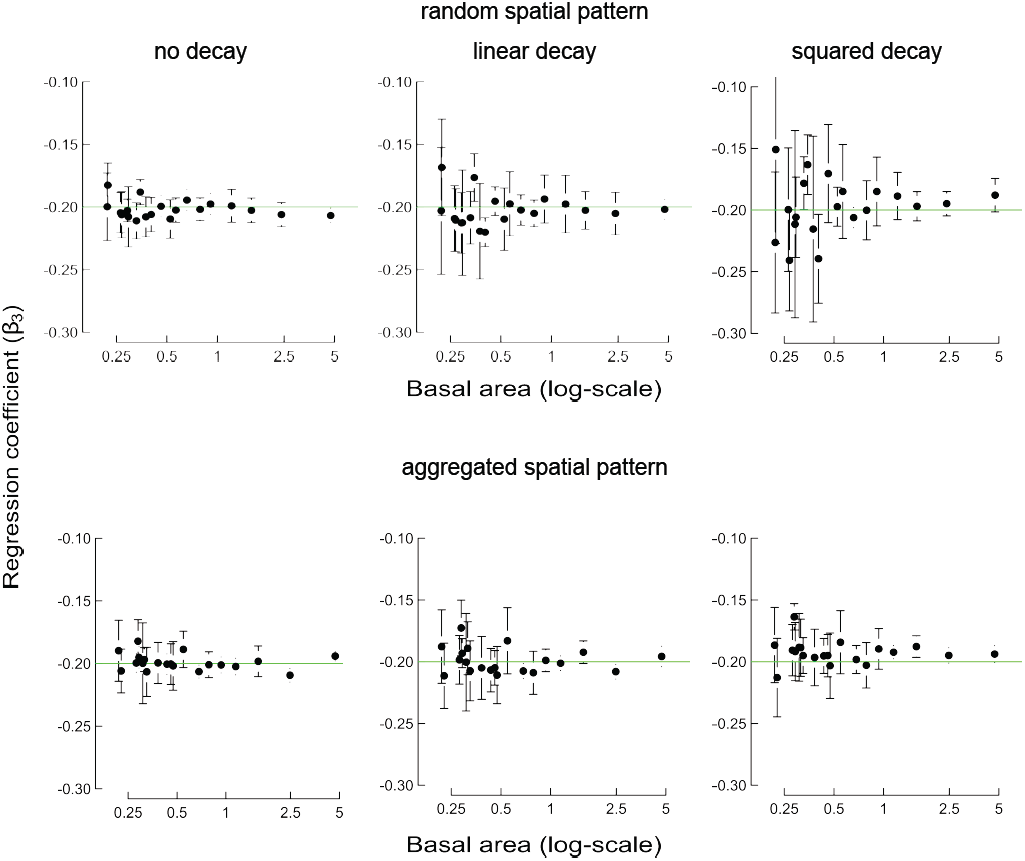
Regressions of conspecific negative density dependence (regression coefficient, β_3_ in Eq. 1) and species abundances (plot level basal area). Twenty species with identical initial size distributions and log-series abundances were simulated without, linear (1/distance) or squared (1/distance^2^) distance decay of conspecific neighbor effects within 20 m radius in random or aggregated spatial patterns. Data points are means (± 1 SD) from five replicate simulations. The simulated input value of β_3_ was -0.2 (green line).

Variance in local conspecific neighbor density (within 20 m) varied depending on distance decay and spatial pattern (Fig. 3). A strong negative regression with abundance emerged without distance decay in both spatial patterns. With linear distance decay, the regression was not significant with random spatial pattern but still negative in the aggregated pattern. With squared distance decay, the regression switched to positive in the random pattern, but it was not significant in the aggregated pattern.

**Figure. 3.**
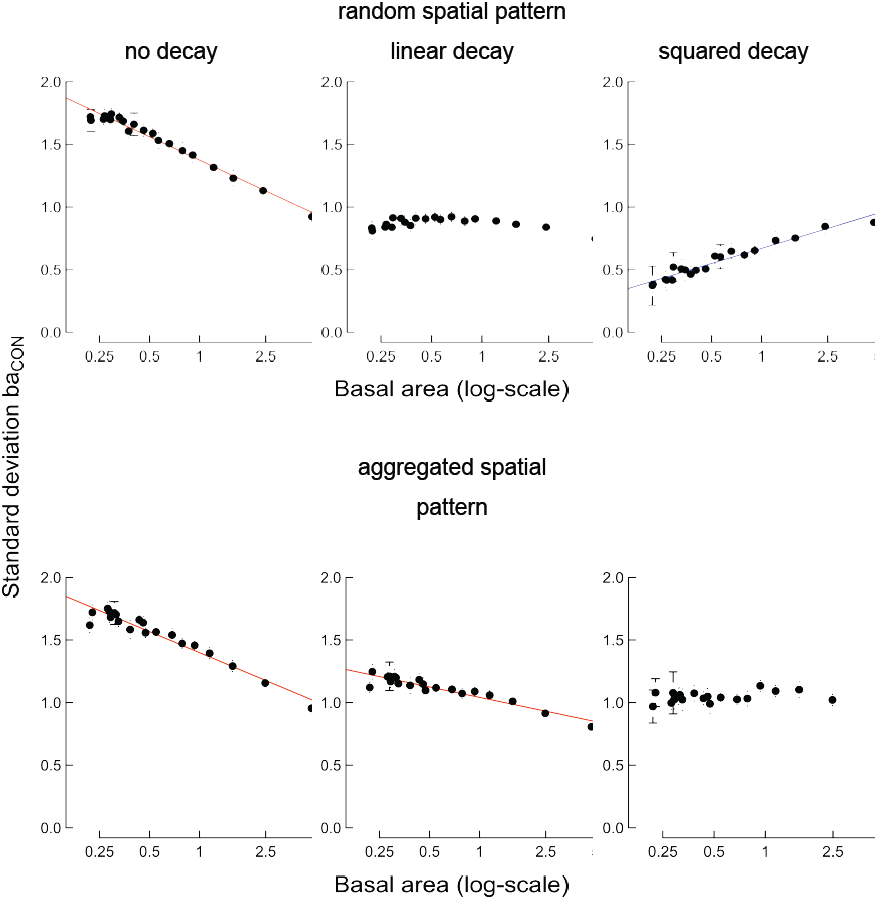
Regressions of variation in conspecific neighbor density (expressed as SD in basal area of bigger conspecifics, ba_CON_ within 20 m) and species abundances (plot level basal area). Twenty species with identical initial size distributions and log-series abundances were simulated with random or aggregated spatial patterns without, linear (1/distance) or squared (1/distance^2^) distance decay of conspecific neighbor effects. Data points are means (± 1 SD) from five replicate simulations. Continuous lines indicate significant (P < 0.05) negative (red) or positive (blue) regressions.

As a consequence of variation in local conspecific neighbor density, effect sizes (Fig. 4) and standardized regression coefficients (b_3_, Fig. 5) showed various relations with abundance depending on distance decay and spatial pattern. Without distance decay both effect sizes and standardized regression coefficients were positively related with abundance, regardless of spatial pattern. This was also the case for effect sizes and linear distance decay, whereas standardized regression coefficients were not significantly related with abundance in random spatial pattern but still positively related with abundance in the aggregated pattern. For squared distance decay, both effect sizes and standardized regression coefficients were negatively related with abundance in random spatial patterns but unrelated in aggregated patterns. Apparently, the squared distance decay canceled the effect of aggregation.

**Figure. 4.**
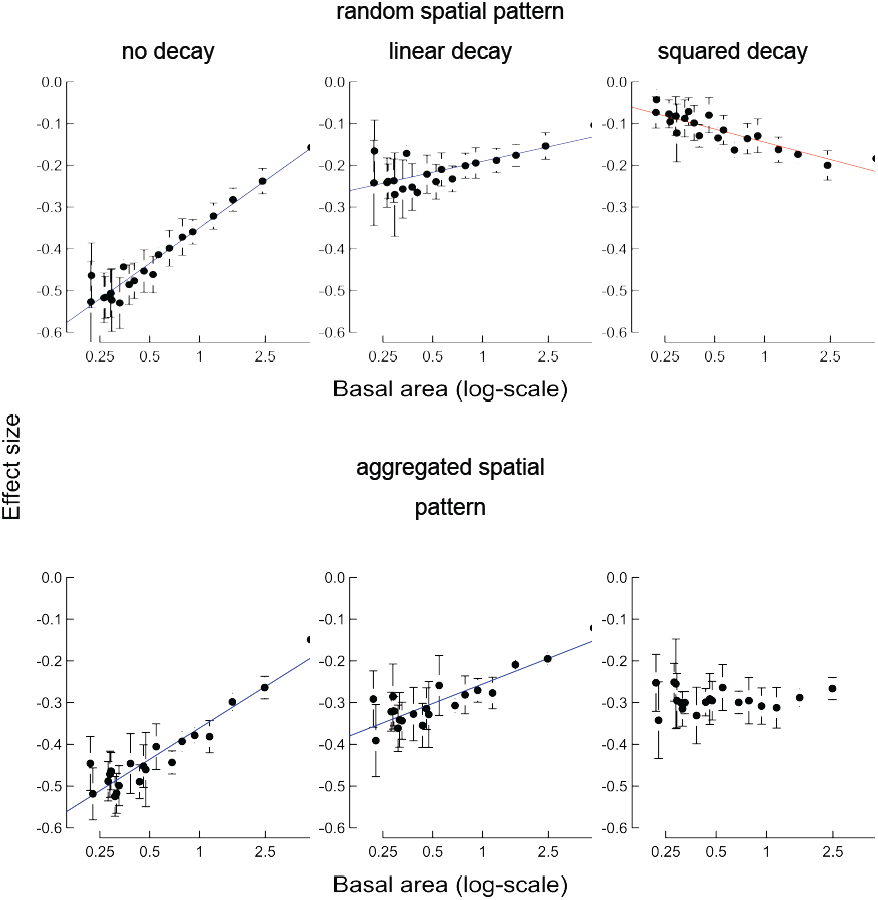
Regressions of effect sizes (squared partial correlation coefficients of β_3_ in Eq. 1) and species abundances (plot level basal area). Twenty species with identical initial size distributions and log-series abundances were simulated with random or aggregated spatial patterns without, linear (1/distance) or squared (1/distance^2^) distance decay of conspecific neighbor effects within 20 m radius. Data points are means (± 1 SD) from five replicate simulations. Continuous lines indicate significant (P < 0.05) positive (blue) or negative (red) regressions.

**Figure. 5.**
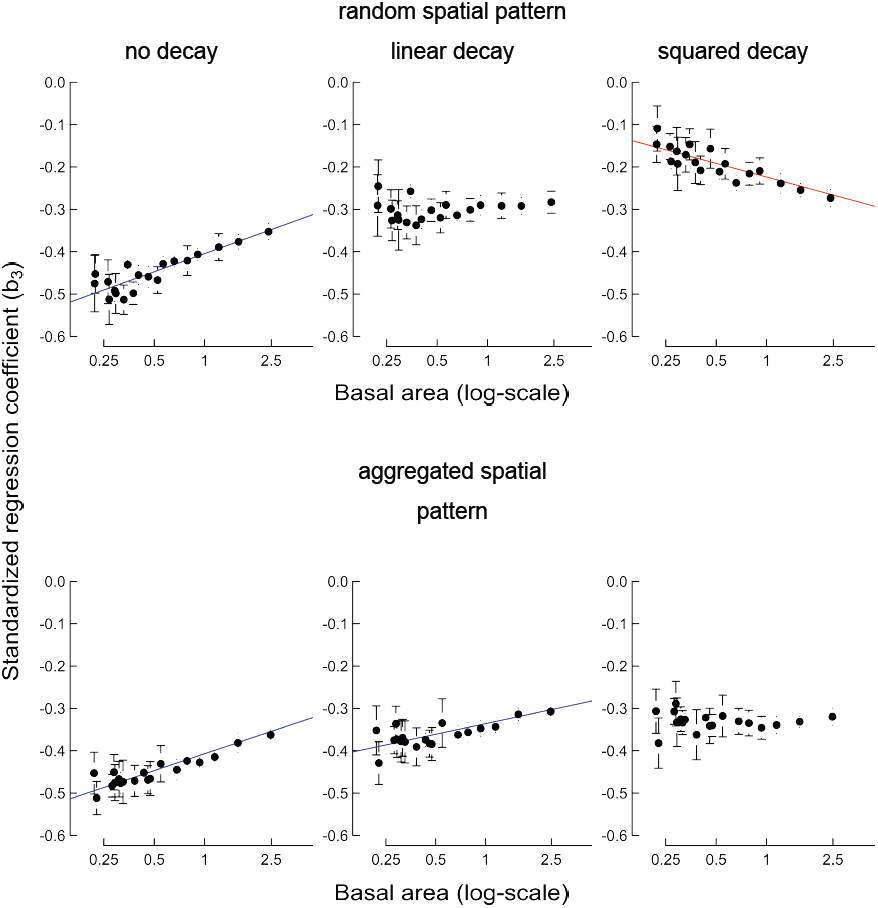
Regressions of standardized regression coefficients (b_3_) and species abundances (plot level basal area). Twenty species with identical initial size distributions and log-series abundances were simulated with random or aggregated spatial patterns without, linear (1/distance) or squared (1/distance^2^) distance decay of conspecific neighbor effects within 20 m radius. Data points are means (± 1 SD) from five replicate simulations. Continuous lines indicate significant (P < 0.05) positive (blue) or negative (red) regressions.

## 4 Discussion

Our simulations and neighborhood analyses showed that estimates of regression coefficients were unrelated to species abundances independent of spatial pattern and distance decay — as expected based on the simulations of identical species without species-specific interactions. However, variation in local density of conspecifics showed various relationships with species abundances depending on degree of spatial pattern and form of distance decay. As a consequence, relationships between effect sizes, or standardized regression coefficients, and species abundances were either non-significant, positive or negative.

By using neighborhood models without distance decay and unstandardized input variables, in single-species analyses, we found a negative relationship between CNDD and forest-level abundance, at least for the first of the two 10-year periods analyzed (Newbery and Stoll 2013). Using no distance decay, yet standardizing before fitting their models, Lin et al. (2012) found positive relationships over their dry-season interval. Using an exponential distance decay, Comita et al. (2010) centered (subtracted the mean) but did not standardize (divide by standard deviation) their input variables (L. Comita, pers. comm.) and found a strong positive relationship too. Whereas Lin et al. (2012) fitted mixed models using maximum likelihood estimation, i.e. without any prior information being involved, Comita et al. (2010) used a hierarchical Bayesian analysis with non-informative priors distributed according to the scaled inverse-Wishart function. This conjugate distribution models the covariance matrix of the species-level regression. Nevertheless, both studies did find positive relationships, thereby apparently supporting one another’s conclusions.

We wondered, though, whether the specific scale and distribution of the priors used by Comita et al. (2010) might have introduced additional critical information that determined in part the estimation of their coefficients, in a similar way as standardization did in our simulations, and may also have done for Lin et al. (2012). Gelman and Hill (2007), for example, discuss the use of the inverse Wishart distribution in some detail. They highlight in particular the need to confirm that Bayesian priors were indeed non-informative across the same ranges of independent variables that resulted in the posterior probabilities. Similarly, Dennis (1996) discussed more general and fundamental issues concerning the use of non-informative priors and Bayesian analysis in ecology.

We are aware that our results dealt with effects of conspecific neighbors (as large tree abundance) on the growth of small trees, whereas those of Comita et al. (2010) concerned conspecific neighbor effects (as either seedling density or tree abundance) on survival of seedlings. Their interesting result would be more generally important if it could be shown to be fully robust. It might then contribute to the notion (Uriarte et al. 2004 ab, Newbery and Stoll 2013) that fundamentally different DD processes are most likely operating at the seedling as opposed to the small tree stage in tropical forest dynamics.

Because of conceptual similarities of neighborhood analyses of Comita et al. (2010) and those of others (e.g., Lin et al. 2012; Uriarte et al. 2004ab), our analysis here could be more widely relevant. Since standardization can lead to spurious relationships between CNDD and species abundances — as we have shown here, its potential influence needs to be carefully considered when interpreting relationships of small-scale effects of conspecific neighbors on larger scale abundance patterns within diverse tree communities. Similarly, care should be taken when specifying prior information in hierarchical Bayesian analyses. Our recommendation, following from Newbery and Stoll (2013), is that tests that randomize tree positions and identities indeed provide the best benchmark by which to critically evaluate and judge relationships between effect sizes, or standardized regression coefficients, and tree species abundances.

## 5 Supplementary Materials

### Appendix A

Documented computer code used for the simulations. A detailed description of input parameters and simulation output is provided in the file named growth_files.rtf.

## References

Bagchi R et al. (2014) Pathogens and insect herbivores drive rainforest plant diversity and composition. Nature 506:88

Cohen J (1988) Statistical power analysis for the behavioral sciences, Second edn. L. Earlbaum Associates, USA

Comita LS, Muller-Landau HC, Aguilar S, Hubbell SP (2010) Asymmetric density dependence shapes species abundances in a tropical tree community. Science 329:330–332. doi: 10.1126/science.1190772

Connell JH, Tracey JG, Webb LJ (1984) Compensatory recruitment, growth, and mortality as factors maintaining rain forest tree diversity. Ecological Monographs 54:141–164

Dennis B (1996) Discussion: Should ecologists become Bayesians? Ecological Applications 6:1095–1103. doi: 10.2307/2269594

Gelman A, Hill J (2007) Data analysis using regression and multilevel/hierarchical models. Cambridge University Press, Cambridge

Lin LX, Comita LS, Zheng Z, Cao M (2012) Seasonal differentiation in density-dependent seedling survival in a tropical rain forest. Journal of Ecology 100:905–914. doi: 18 10.1111/j.1365-2745.2012.01964.x

Muller-Landau HC (2014) Ecology: Plant diversity rooted in pathogens. Nature 506:45

Nakagawa S, Cuthill IC (2007) Effect size, confidence interval and statistical significance: a practical guide for biologists. Biological Reviews 82:591–605. doi: 10.1111/j.1469-22 185X.2007.00027.x

Newbery DM, Stoll P (2013) Relaxation of species-specific neighborhood effects in Bornean rain forest under climatic perturbation. Ecology 94:2838–2851. doi: 10.1890/13-0366.1

Neyman J, Scott EL (1952) A theory of the spatial distribution of galaxies. Astrophysical Journal 116:144–163. doi: 10.1086/145599

R Development Core Team (2012) R: A language and environment for statistical computing. R Foundation for Statistical Computing, Vienna, Austria

Schielzeth H (2010) Simple means to improve the interpretability of regression coefficients. Methods in Ecology and Evolution 1:103–113. doi: 10.1111/j.2041-210X.2010.00012.x

Stoll P, Newbery DM (2005) Evidence of species-specific neighborhood effects in the K dipterocarpaceae of a Bornean rain forest. Ecology 86:3048–3062. doi: 10.1890/04-1540

Uriarte M, Condit R, Canham CD, Hubbell SP (2004a) A spatially explicit model of sapling growth in a tropical forest: does the identity of neighbours matter? Journal of Ecology 92:348–360

Uriarte U, Canham CD, Thompson J, Zimmerman JK (2004b) A neighborhood analysis of tree growth and survival in a hurricane-driven tropical forest. Ecological Monographs 14 74:591–614

Warner RM (2012) Applied statistics: from bivariate through multivariate techniques. Sage, Los Angeles, USA

